## Supplemental for "ER O-glycosylation in synovial fibroblasts drives cartilage degradation"

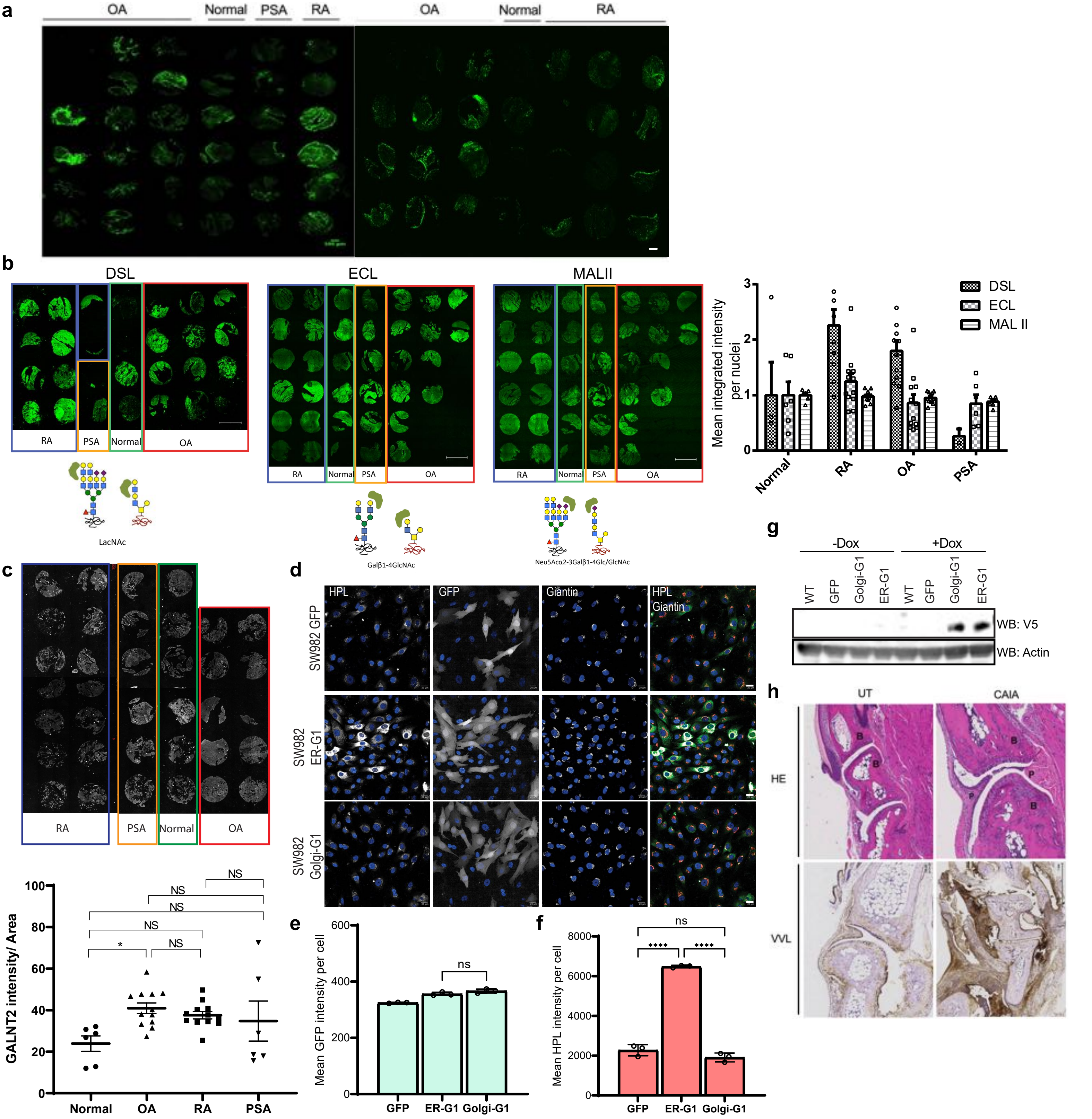


Supplementary Fig. 1: O-GalNAc (Tn) glycans tissue analysis in synovial tissues from both arthritis patients and arthritis induced animals
(a) Representative immunofluorescence staining of O-GalNAc glycans with VVL lectin on human tissue microarray (TMA) sections from Osteoarthritis (OA), psoriasis (PSA), rheumatoid arthritis (RA) and health subjects (Normal). Magnification 10X, scale bar, 500 µm. (b) Immunofluorescence staining of human tissue array with lectins DSL (Datura Stramonium Lectin), ECL (Erythrina Cristagalli Lectin) and MAL II (Maackia Amurensis Lectin). Glycan strucures targeted by the lectins were shown below each panel. Quantification of the lectin staining intensities shown on the right. (c) Immunofluorescence staining of human tissue array with GALNT2 antibody. Quantification of the staining intensities shown below. (d) Representative images of Tn staining using HPL in SW982 cell lines expressing control GFP, ER-localised GALNT1 (ER-G1) and wildtype GALNT1 (Golgi-G1).Scale bar, 20 µm. (e) Quantification of total GFP fluoresence intensity in control GFP, ER-G1-GFP and Golgi-G1-GFP expressing SW982 cells. NS, not significant (one-way ANOVA test). (f) Quantification of HPL intensity in GFP, ER-G1 and Golgi-G1 expressing SW982 cells. Data are shown as mean ± SD. ****, p<0.0001, NS, not significant (one way ANOVA test). (g) Levels of exogenously expressed GALNT1 in wildtype SW982 (WT), control GFP (GFP), wildtype GALNT1 (Golgi-G1) and ER-localised GALNT1 (ER-G1) that were uninduced (-Dox) or induced with 1µg/ml doxycycline (+Dox) over 24hours. Exogenously expressed GALNT1 have a V5 tag. (h) HE histology (upper panel) and immunohistochemistry staining with VVL lectin (lower panel) on collagen type II antibody induced arthritis (CAIA) mice at day 7 or those left untreated (UT).

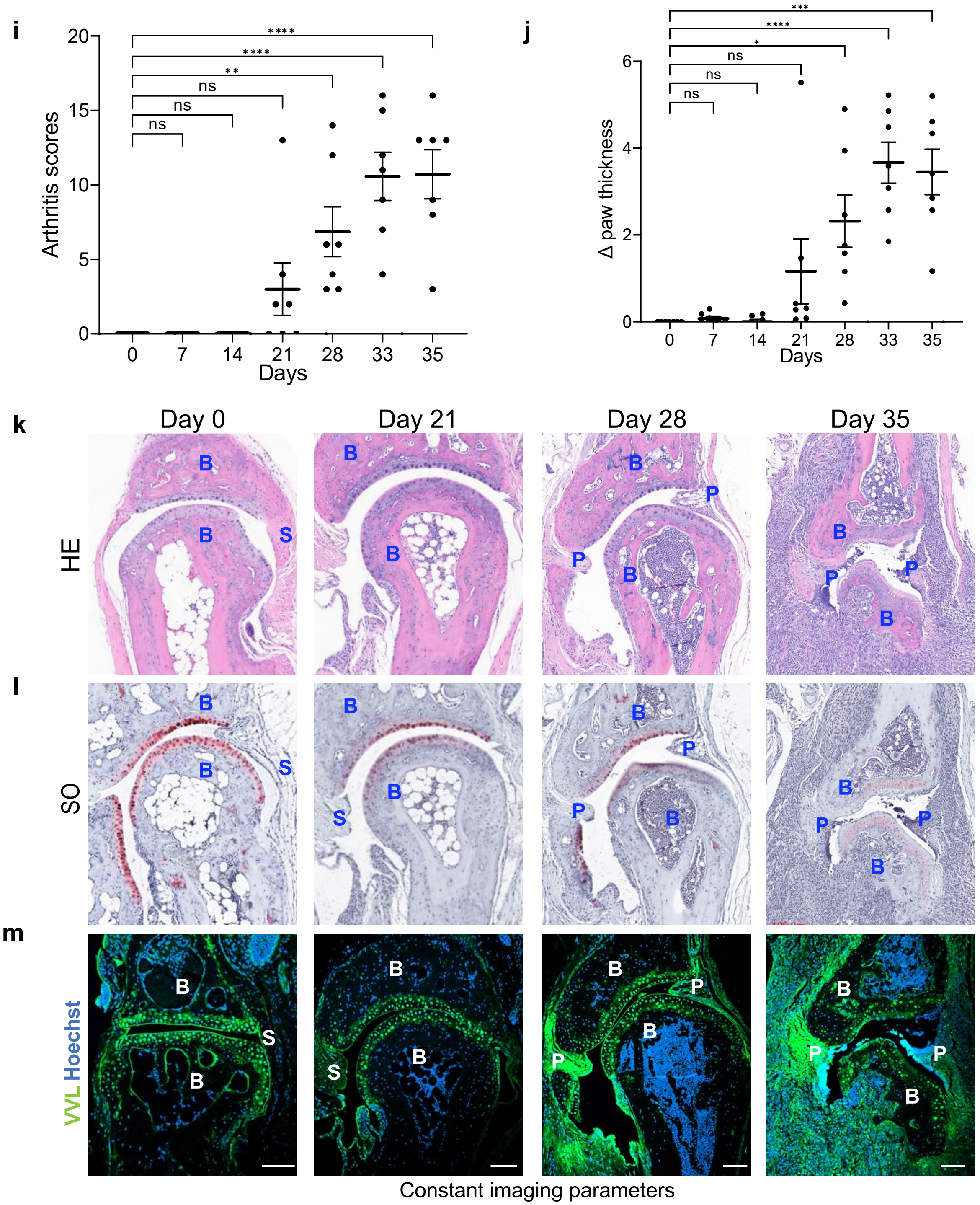


Supplementary Fig. 1: O-GalNAc (Tn) glycans tissue analysis in synovial tissues from both arthritis patients and arthritis induced animals
(i) Clinical scores of CIA mice from day 0 to day 35. Mean ± SD of the arthritic scores of 7 mice per time point presented. (j) Measurement of the change in paw thickness in CIA mice from day 0 to day 35. Mean ± SD of 7 mice per time point presented. (k) Hematoxylin and eosin (HE) histology images of synovial tissues sections from control (day 0) or CIA mice at day 21, 28 and 35. S: synovium; B: bone, P: pannus. (l) Safranin O (SO) histology images of synovial tissues sections from control (day 0) or CIA mice at day 21, 28 and 35. (m) VVL lectin (green) and nuclei (Hoescht) images of synovial tissues sections from control (day 0) or CIA mice at day 21, 28 and 35. Images were acquired under constant parameters. Scale bar, 100 μm. Notice strongly VVL stained panus (P) at day 28 and 35.

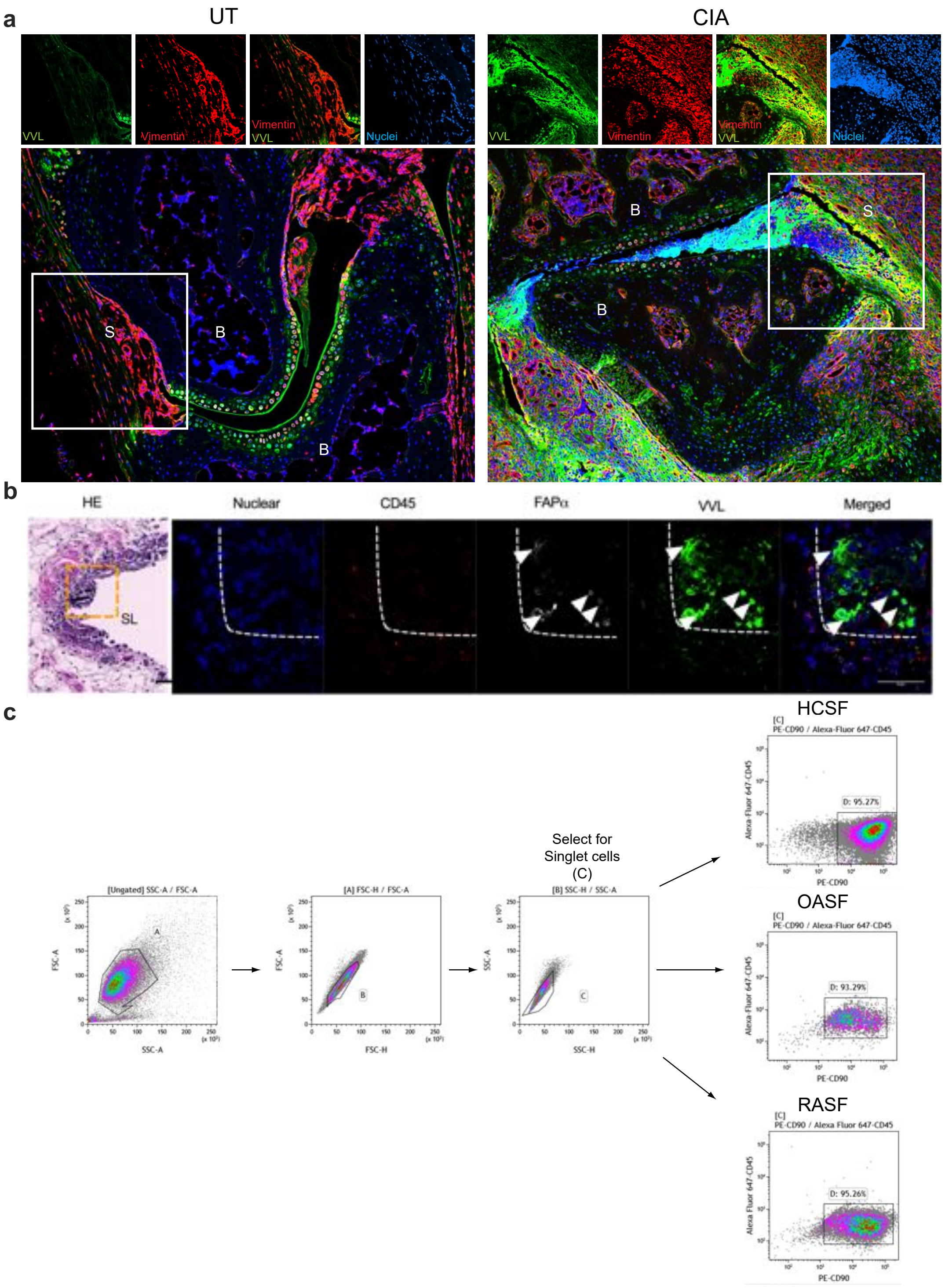


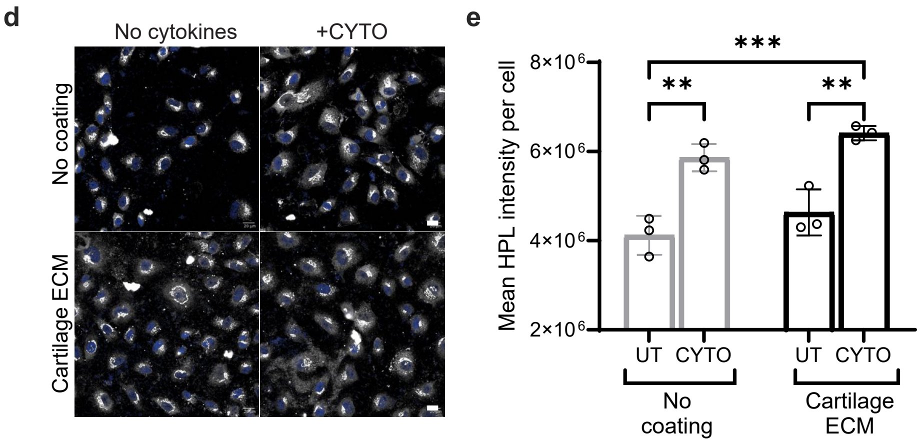


Supplementary Fig. 2: Synovial tissue analysis and primary synovial fibroblasts purification metric. (a) Representative immunofluorescence images of joints from CIA mouse on day 28 post induction. Mouse fibroblasts are stained with fibroblast maker vimentin and nuclei with hoescht staining; Scale bar, 50 µm. S: synovium; B: bone. (b) (left) (H&E) histology of synovial tissues obtained from OA patients. Scale bar, 100 µm; SL: synovial lining. (right) Representative immunofluorescence images of OA synovium stained with VVL/CD45 and FAPa. Lining synovial fibroblasts are identified by FAPa (arrowheads).Immune cells are identified by CD45. Sublining layer and lining layers boundaries are defined with dash line. Scale bar, 50 µm. (c) Gating strategy of the flow cytometry analysis of primary SF cultures from synovial tissues of healthy subjects (HCSF), OA (OASF) and RA (RASF) patients. Dot plots showing high purity (>90%) of fibroblast cells. (d) Representative images of wildtype SW982 cells stimulated with or without CYTO and cartilage ECM stained with HPL lectin in white. Scale bar: 20 µm. (e) Quantification of HPL intensity per cell in wildtype SW982 under basal conditions (UT) and upon presence of cytokines IL1β and TNFα (CYTO) plated on plastic (No coating) or plated on cartilage ECM. Data are mean ± SD of 3 replicates. *, p < 0.05, **, p< 0.01, ***, p< 0.001, (Two-way ANOVA).

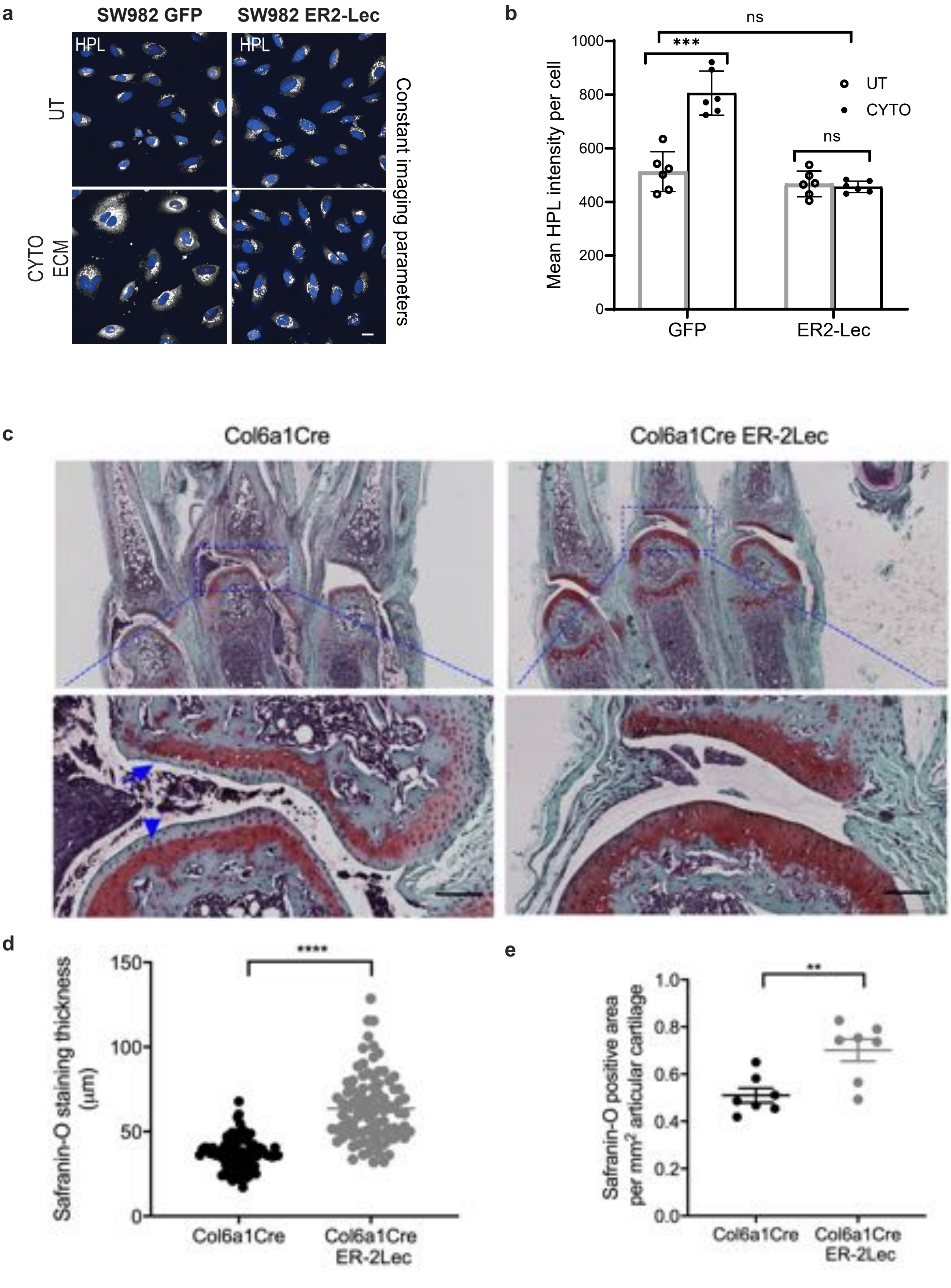


Supplementary Fig. 3: GALA activation causes damage to cartilage in CAIA mice. (a) Tn staining in SW982 cells expressing GFP or ER2-Lec. Scale bar, 20 μ m. (b) Quantification of Tn levels in (a). Data are mean ± SEM of two 2 independent experiments. *, $p < 0.05$, ***, $p< 0.001$, ****, $p< 0.0001$, NS: not significant (One way ANOVA). (c) Representative images of Safranin-O (SO) (B) staining in untreated Col6a1Cre mice or arthritis induced Col6a1Cre and Col6a1Cre ER-2Lec mice at day 7. Scale bars, 100 μm. (d & e) Quantification of SO staining thickness (d) and total positive staining area (e). Arthritis induced cartilage matrix degradation is indicated with arrowheads . Each data point represents an average of positive staining area per mm2 articular cartilage from three different metacarpophalangeal joints of one animal. Data are shown as mean ± SEM. $p<0.01$; ****, $p<0.0001$ (one way ANOVA test).

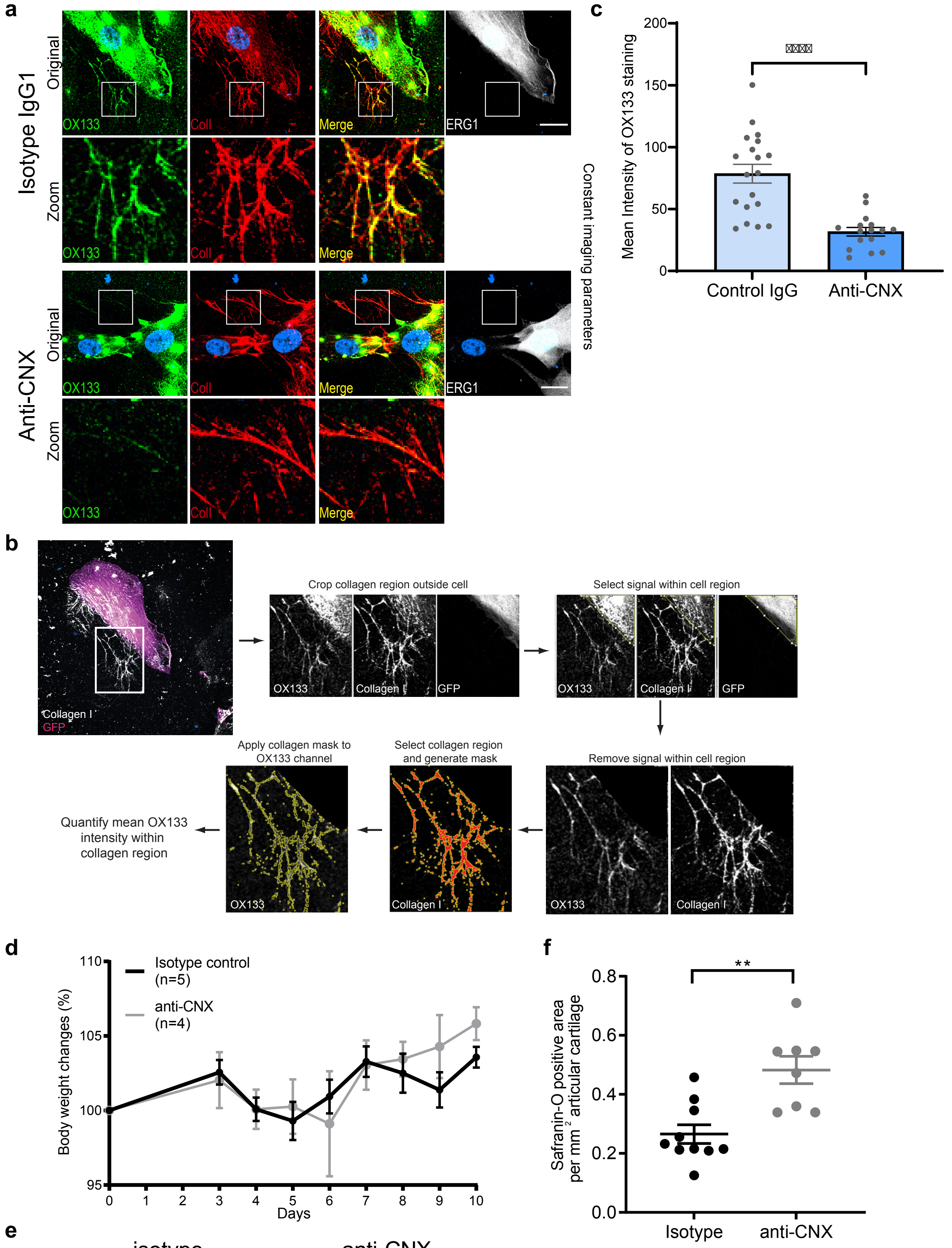


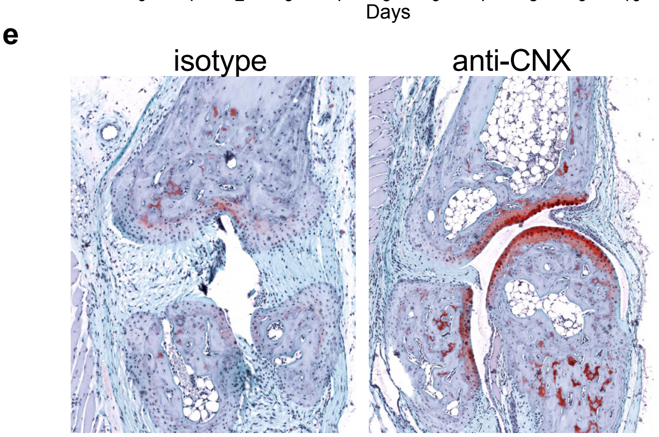


Supplementary Fig. 4: Treatment with antibodies against CNX protects CAIA mice from cartilage degradation
(a) Representative images of OX133 staining on Collagen I fibres of cartilage ECM around SW982 ER-G1 cells in the presence of isotype IgG1 antibody or anti-CNX antibody. Scale bar, 20um.
(b) ImageJ analysis workflow to quantify OX133 staining intensities on collagen I fibres.
(c) Quantification of OX133 staining intensities on collagen I fibres around cells that were treated with either isotype IgG1 antibody or anti-CNX antibody . Data are shown as mean ± SEM ****, p<0.0001 (one way ANOVA test).
(d) Body weight changes of CAIA mice treated with isotype control or anti-CNX antibodies
(e & f) Representative histological images (e) and quantification (f) of Safranin-O (SO) staining areas in CAIA mice treated with isotype control or anti-CNX antibodies. Individual data points represent average of positive staining area per mm2 articular cartilage from individual metacarpophalangeal joints of one animal, n= 4-5 mice per group. Data are shown as mean ± SEM. *, p<0.05; **, p<0.01(Mann-Whitney test).

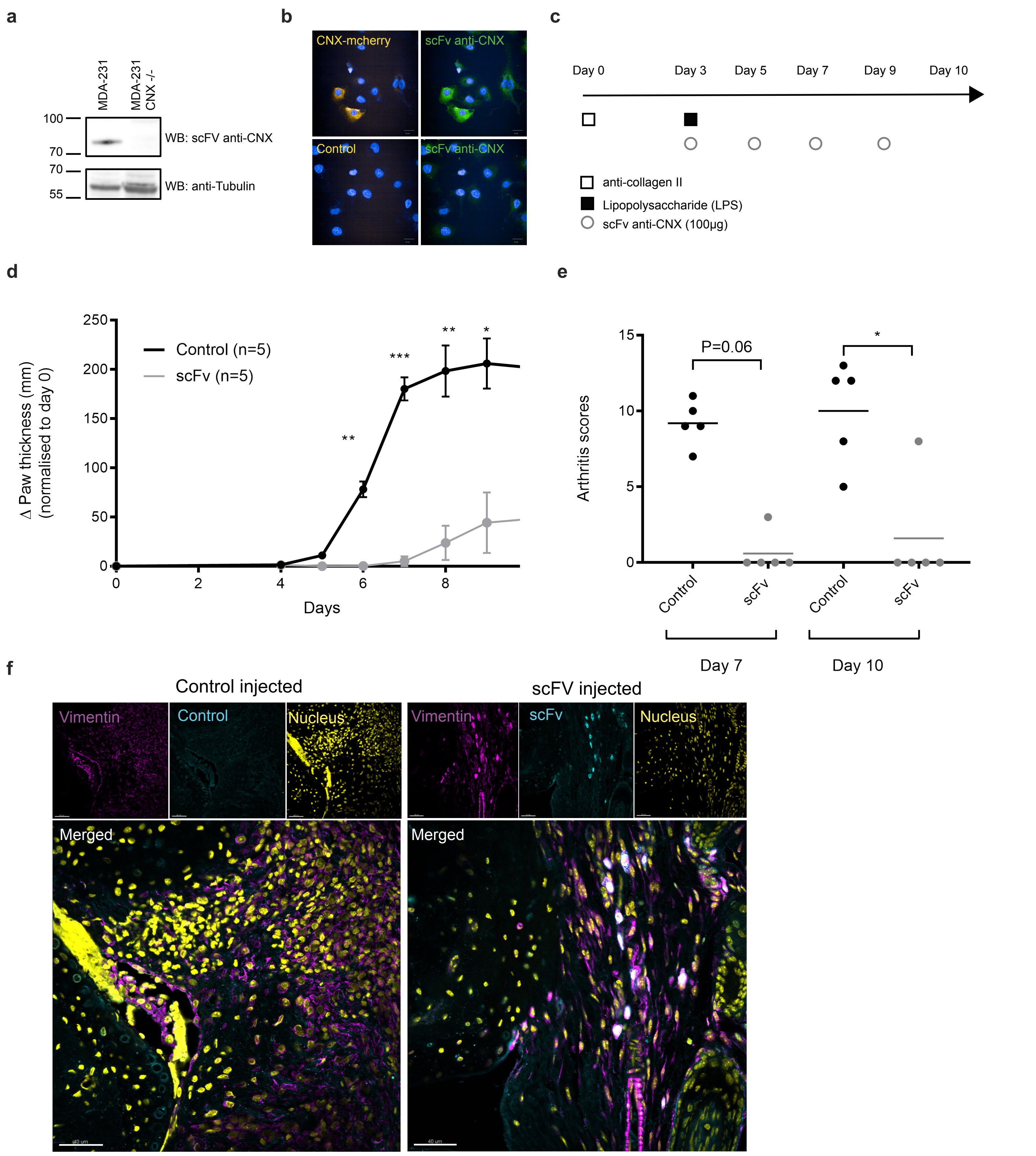


Supplementary Fig. 5: Blockage of CNX with single chain Fv (scFv) protects CAIA mice from arthritis.
(a) Western Blot analysis of CNX with identified scFv against CNX. Cell lysates from MDA-231 and MDA-231 CNX-/- probed with myc tagged scFv specific for CNX and subsequently with anti-myc Horseradish peroxidase. A loading control with tubulin is indicated. (b) Representative immunofluorescence images with scFv against CNX. Fixed and permeabilized Huh7 normal cells or Huh7 cells with stably integrated mcherry CNX (orange) subjected to incubation with myc tagged scFv and subsequently to secondary anti-Myc conjuguated to PhycoErythrin (green) and Hoechst nuclear dye (blue). Representative pictures for CNX mcherry stable cells are indicated on top (left) along with scFv co-stain (right). Representative pictures for Huh7 control cells are indicated on the bottom (left) along with scFv co-stain (right). All images are from constant acquisition and display settings. scale bar 20μm. (c) Experimental setup of injections and timeline used for treatment of collagen antibody induced arthritis mouse model with scFv against CNX. (d) Paw thickness variation in CAIA mice treated with scFv injection or control PBS injection from day 0 to day 10. (e) Clinical scores of CAIA mice treated with scFv injection or control PBS injection from day 0 to day 10. (f) Representative immunofluorescence images of synovial tissues sections from CAIA mice day 10 from control PBS or myc tagged scFv injected mice stained with anti-Myc conjuguated to PhycoErythrin (green), Vimentin and DAPI. Scale bar, 40 μm.
